# *Burkholderia pseudomallei* BicA protein promotes pathogenicity in macrophages by regulating Invasion, Intracellular Survival, and Virulence

**DOI:** 10.1101/2023.07.10.548404

**Authors:** Jacob L Stockton, Nittaya Khakhum, Heather L. Stevenson, Alfredo G Torres

## Abstract

*Burkholderia pseudomallei (Bpm)* is the causative agent of melioidosis disease. *Bpm* is a facultative intracellular pathogen with a complex lifecycle inside host cells. Pathogenic success depends on a variety of virulence factors with one of the most critical being the type 6 secretion system (T6SS). *Bpm* uses the T6SS to move into neighboring cells, resulting in multinucleated giant cells (MNGCs) formation, a strategy used to disseminate from cell-to-cell. Our prior study using a dual RNA-seq analysis to dissect T6SS-mediated virulence on intestinal epithelial cells identified BicA as a factor upregulated in a T6SS mutant (1). BicA regulates both type 3 secretion system (T3SS) and T6SSs; however, the extent of its involvement during disease progression is unclear. To fully dissect the role of BicA during systemic infection, we used two macrophage cell lines paired with a pulmonary *in vivo* challenge murine model. We found that Δ*bicA* has a distinct intracellular replication defect in both immortalized and primary macrophages that begins as early as 1 h post-infection. This intracellular defect is linked with the lack of cell-to-cell dissemination and MNGC formation, as well as a defect on T3SS expression. The *in vitro* phenotype translated *in vivo* as Δ*bicA* was attenuated in a pulmonary model of infection; demonstrating a distinct macrophage activation profile and lack of pathological features present in the wild type. Overall, these results highlight the role of BicA in regulating intracellular virulence and demonstrate that specific regulation of secretion systems has a significant effect on host response and *Bpm* pathogenesis.

**Importance:** Melioidosis is an understudied tropical disease that still results in ∼50% fatalities from those infected patients. It is caused by the Gram-negative bacillus *Burkholderia pseudomallei* (*Bpm*)*. Bpm* is an intracellular pathogen that disseminates from the infected cell to target organs, causing disseminated disease. Regulation of secretion systems involved in entry and cell-to-cell spread is poorly understood. In this work, we characterize the role of BicA as a regulator of secretion systems during infection of macrophages *in vitro* and *in vivo*. Understanding how these virulence factors are controlled will help us determine their influence on the host cells and define the macrophage responses associated with bacterial clearance.

## Introduction

*Burkholderia pseudomallei (Bpm)* is a Gram-negative, environmental saprophyte that causes the disease melioidosis (2). Melioidosis is a neglected tropical disease with an estimated 165,000 cases per year with 89,000 fatalities; however, these numbers seem to be underreported due to identification of new endemic areas. It was thought to be restricted to southeast Asia and Australia but it has become clear that it is present in some capacity in most tropical and sub-tropical regions of the globe (3). This includes the United States where *Bpm* has been routinely imported (4, 5) but evidence is beginning to suggest endemicity in the Gulf Coast regions, particularly Mississippi (6, 7). Combating *Bpm* is particularly difficult due to the lack of a licensed vaccine (8), extensive antibiotic resistance, ability to generate persistent infections (9, 10), and its intracellular lifestyle (2). As a facultative intracellular pathogen, *Bpm* utilizes a myriad of virulence factors to promote replication including the type VI secretion system (T6SS) (11). The T6SS is a contractile nanomachine widely distributed across Gram-negative species that is primarily used to deliver effector proteins for interbacterial competition (12). There is a small subset of T6SSs that have eukaryotic targets, including *Bpm* which uses the T6SS to spread from cell-to-cell via cell fusion events, resulting in multinucleated giant cell (MNGC) formation (11, 13). MNGC formation is the keystone event during the intracellular pathogenesis process but the mechanisms behind the membrane fusion events are largely unknown, from both the bacterial and host sides.

Recently, we began investigating the mechanisms of T6SS-mediated virulence within the context of the understudied gastrointestinal (GI) route of infection (14) and performed a dual RNA-seq analysis with primary murine intestinal epithelial cells (pMIECs) infected with wild-type (WT) *Bpm* or a T6SS structural mutant *Δhcp1* (BPSS1498) (1). In this analysis, *bicA* (BPSS1533) was identified as significantly upregulated in *Δhcp1*, and this was particularly interesting as BicA has been implicated in the regulatory network controlling both type III secretion system (T3SS) and T6SS expression (15). It has been suggested that BicA is the chaperone/co-activator of BsaN and together they act to coordinate the timely expression of T3SS effectors and downstream T6SS regulator genes *bprC* and *virAG* (16). Interestingly, *bsaN* was significantly downregulated in *Δhcp1,* which suggests a different involvement of BicA in the regulation of T3SS and T6SS. We evaluated the contribution of BicA within our GI model of infection and determined that it is critical for intracellular replication, cell-to-cell spread, and lethality (1). As the dynamics of the GI model of infection are unclear, we began interrogating the mechanisms of *ΔbicA* in the well characterized and systemic macrophage model. Macrophages are multi-function immune cells that are present as specialized tissue resident cells or circulating undifferentiated monocytes. They are capable of a variety of jobs including pathogen clearance, antigen presentation, and immune coordination via secretion of cytokines and chemokines, which greatly influence the inflammatory landscape (17). This makes macrophages an attractive target for subversion by intracellular pathogens like *Bpm,* which uses macrophages as a replicative niche and a dissemination “trojan horse” (18, 19). It has been hypothesized that *Bpm* modulates the activation state of macrophages to promote replication versus clearance, but the mechanism of this modulation is unclear (20, 21).

In this work we demonstrate that BicA is necessary for successful replication inside macrophages using both immortalized and primary macrophages and this intracellular defect starts upon entry into the cell. Further, we characterize the expression profile of critical virulence factors in *ΔbicA* versus WT *Bpm* strains and found a distinct dysregulation of multiple systems. Finally, we examined the role of BicA during inhalational melioidosis and the phenotype of pulmonary macrophages in response to infection. Collectively, we provide evidence to establish BicA as a major regulator of virulence and its absence differentially activates macrophages and leads to clearance of the bacteria.

## Results

### BicA Required for Proficient Intracellular Replication in Macrophages

We previously reported that *ΔbicA* demonstrated a profound intracellular replication defect in intestinal epithelial cells (IECs) (1) so we began our investigation into the role of BicA in the macrophage model by examining intracellular replication in two *in vitro* models: RAW 264.7 (RAW) cells and primary bone marrow-derived macrophages (BMDMs). RAW 264.7 cells are an immortalized macrophage-like cell line originally created from BALB/c mice, so we chose to use BALB/c mice as bone marrow donors for the primary BMDM model. We evaluated the intracellular replication of *Bpm* K96243 WT, *ΔbicA,* and *ΔbicA::bicA* at 3, 6, and 12 h post-infection (hpi) in both RAW cells and BMDMs (**Fig 1A** & **B**) and found that *ΔbicA* replicates at a lower rate as early as 3 hpi and that defect persists through 12 hpi. The replication profiles were nearly identical between macrophage models which led us continue the characterization using both cell models. Since we saw decreased replication at the earliest timepoint of 3 hpi, we wanted to ensure this was not due to differential phagocytosis and to look at the very early stages of infection before the bacteria starts to replicate in the cytoplasm. We measured phagocytosis rates between WT, *ΔbicA,* and *ΔbicA::bicA* and found no significant differences after 1 h of internalization (**Fig 1C**). We also looked at intracellular survival 1 h post-phagocytosis, or 2 h after initial contact of the macrophages with the bacterial inoculum and found that *ΔbicA* exhibited a significant decrease in intracellular survival when compared to WT and *ΔbicA::bicA* (**Fig 1D**). Since bacterial enumeration does not tell a complete story, we visualized BMDMs infected with WT, *ΔbicA, ΔbicA::bicA*, or mock infected at 3, 6, and 12 hpi via immunofluorescence (**Fig 2**). The WT and *ΔbicA::bicA* infected cells showed robust replication at 3 hpi, MNGC formation at 6 hpi, and a sharp decrease in cell viability at 12 hpi. The *ΔbicA* infected cells revealed few intracellular bacteria that, qualitatively, seemed static compared to WT and *ΔbicA::bicA.* These cells also showed no MNGC formation and retained mock levels of viability later into the time course. Together, these data suggest that BicA is critical for intracellular replication and progression through the pathogenesis process.

**Figure 1:**
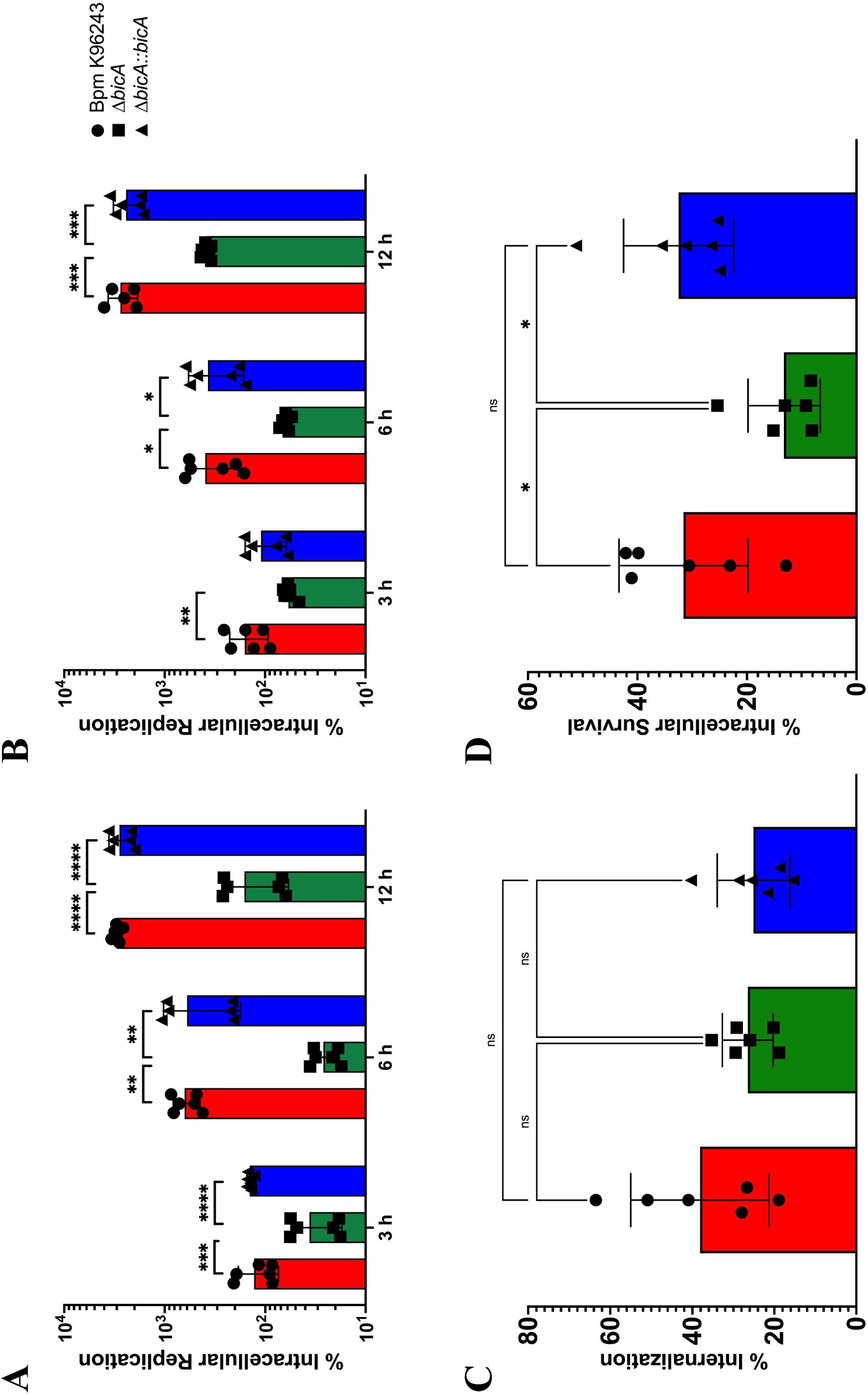
The *ΔbicA* strain demonstrates an intracellular survival defect in macrophages. Macrophages, RAW 264.7 cells (**A**) or BALB/c BMDMs (**B**), were infected at an MOI of 10 with *Bpm* K96243 WT, Δ*bicA,* or Δ*bicA::bicA* and bacteria enumerated at 3, 6, and 12 hpi to assess intracellular replication. Rate of internalization was assessed by incubating bacteria with RAW 264.7 cells for 1 h before enumeration (**C**). Early intracellular survival was assessed by allowing bacteria 1 h for internalization followed by 1 h of 1 mg/mL kanamycin to kill extracellular bacteria (**D**). Bars represent an average of two independent experiments performed in triplicate ± SD. Significant differences were assessed via one-way ANOVA followed by Sidak’s multiple comparison test. p < 0.05 *, p < 0.01**, p < 0.005***, p < 0.0001****.

**Figure 2:**
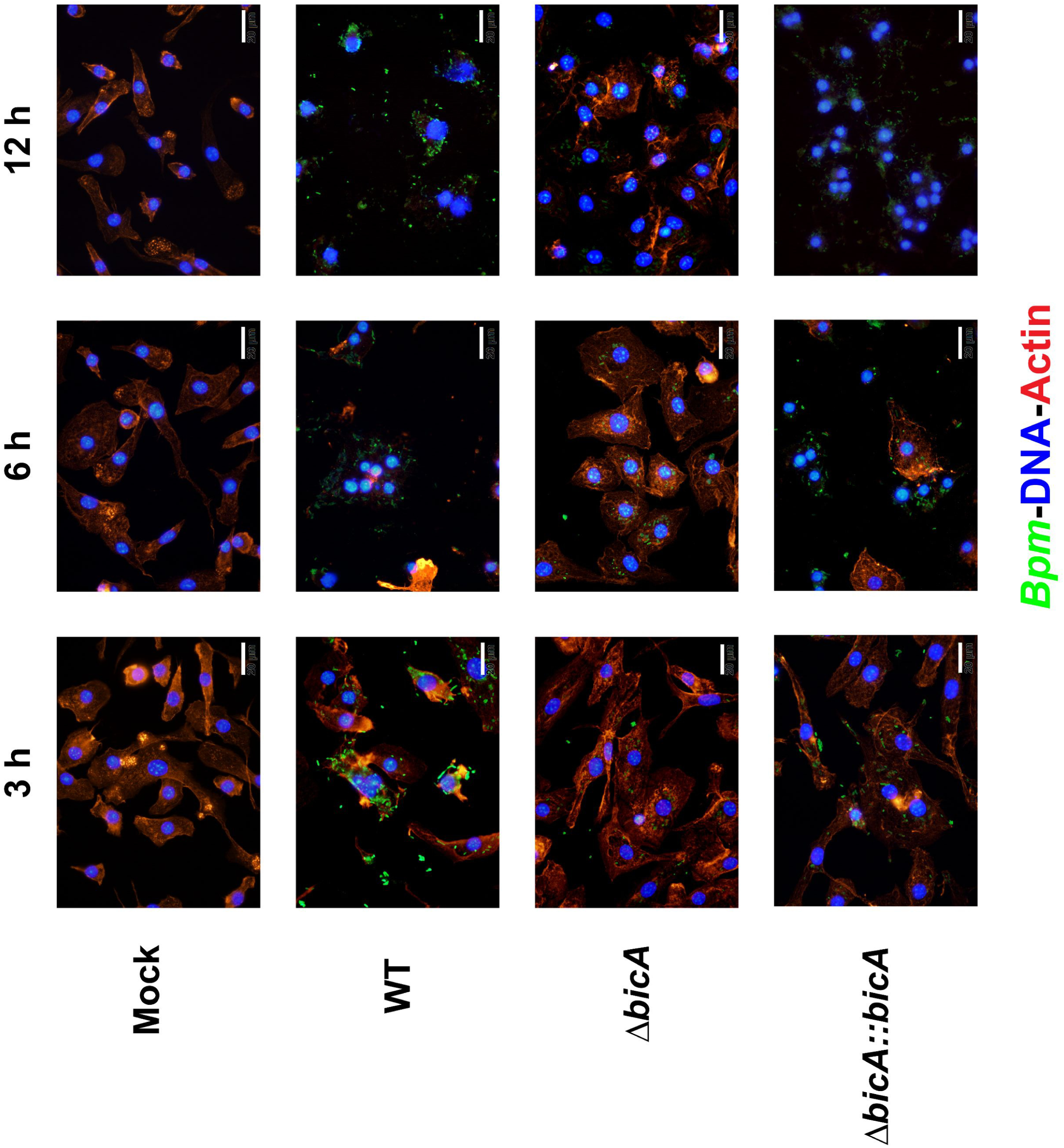
The *ΔbicA* strain appears to exhibit reduced motility and lacks MNGC formation. BALB/c BMDMs were infected at an MOI of 10 with *Bpm* K96243 WT, Δ*bicA,* or Δ*bicA::bicA* for 3, 6, or 12 h before being fixed with 4% PFA and permeabilized with 0.25% Triton-X 100. Bacterial cells were washed and stained with sera from mice vaccinated with a live-attenuated *Bpm* followed by anti-mouse IgG, IgM, IgA (H +L) secondary antibody conjugated to Alexa Fluor 488. Actin and DNA were visualized using rhodamine phalloidin and DAPI, respectively. Images were visualized under 100X magnification using Olympus BX51 upright fluorescence microscope and further analyzed using ImageJ software.

### The *ΔbicA* strain demonstrates disrupted expression of critical virulence factors

Since BicA is predicted to be involved in the regulation cascade of both the T3SS and T6SS machineries, we evaluated the expression of an array of virulence genes during macrophage infection. We selected a panel of genes important during intracellular survival which include the T3SS, T6SS, actin motility proteins, and some of the regulators of these virulence mechanisms (Table 1). RAW 264.7 cells were infected with WT, *ΔbicA,* or *ΔbicA::bicA* and bacterial RNA was extracted at 3 and 6 hpi and semi-quantitative PCR was performed on cDNA libraries generated from bacterial RNA. Beginning at 3 hpi, *ΔbicA* exhibits slight increased expression of T3SS effectors with downregulation of T6SS structural loci and actin motility when compared to WT (**Fig 3**). This trend continues to progress at 6 hpi with high levels of T3SS effector expression and greater downregulation of the T6SS. Actin motility genes appear to be slowly recovering expression as *bimA* is on par with WT at 6 hpi, however, *bimC* is still downregulated. Interestingly, at both 3 and 6 hpi, there appears to be a small increase in activity at the *tssA-virAG* operon but with no evident increase in expression of T6SS structural genes. This operon contains the primary regulators (*virAG*) and structural protein (*tssA*), so the production of these proteins should lead to the production of T6SS machinery but in *ΔbicA* this dynamic appears to be interrupted through an unknown regulatory event. Expression levels of *ΔbicA::bicA* were also compared to WT but no differences were detected (Table S1). The difference in expression of critical virulence factors sheds light on the regulatory impact of BicA and begins to explain the intracellular behavior of *ΔbicA.* The mechanism that restricts the intracellular survival of the Δ*bicA* strain is unknown but it likely tied to alteration of T3SS activity, as this secretion system is responsible for early intracellular success of *Bpm*.

**Table 1.**
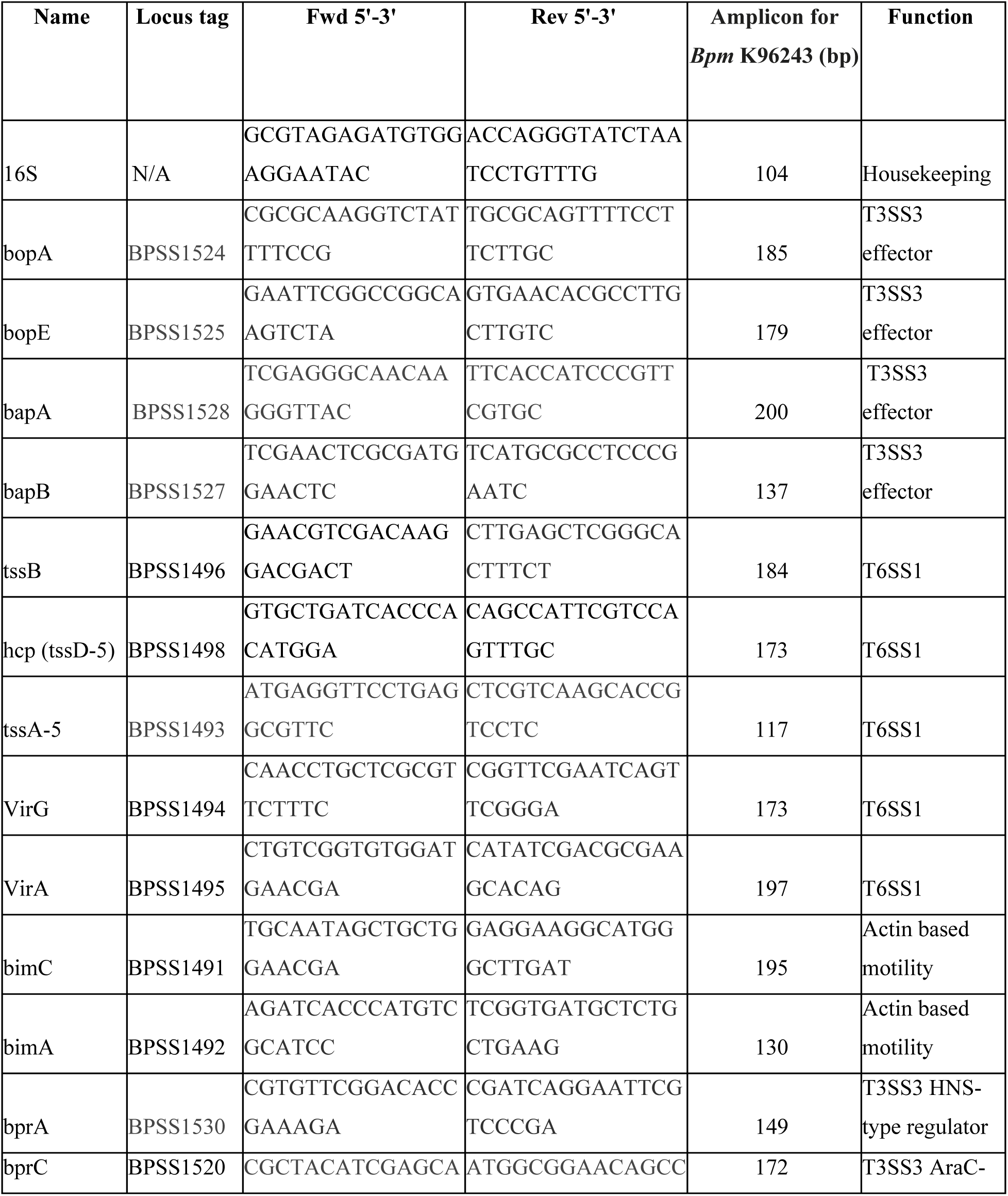
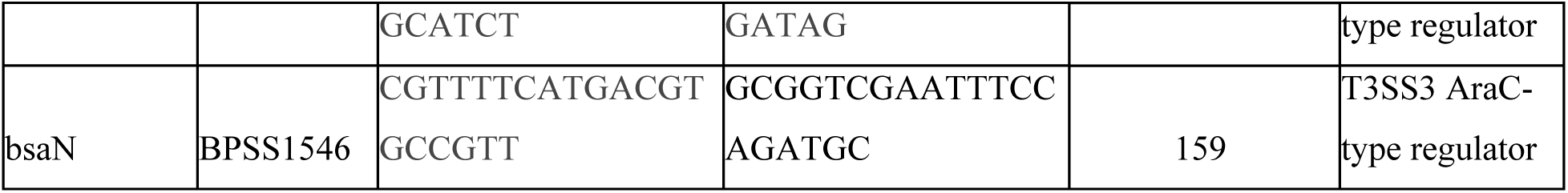
Primers for qPCR used in this study.

**Figure 3:**
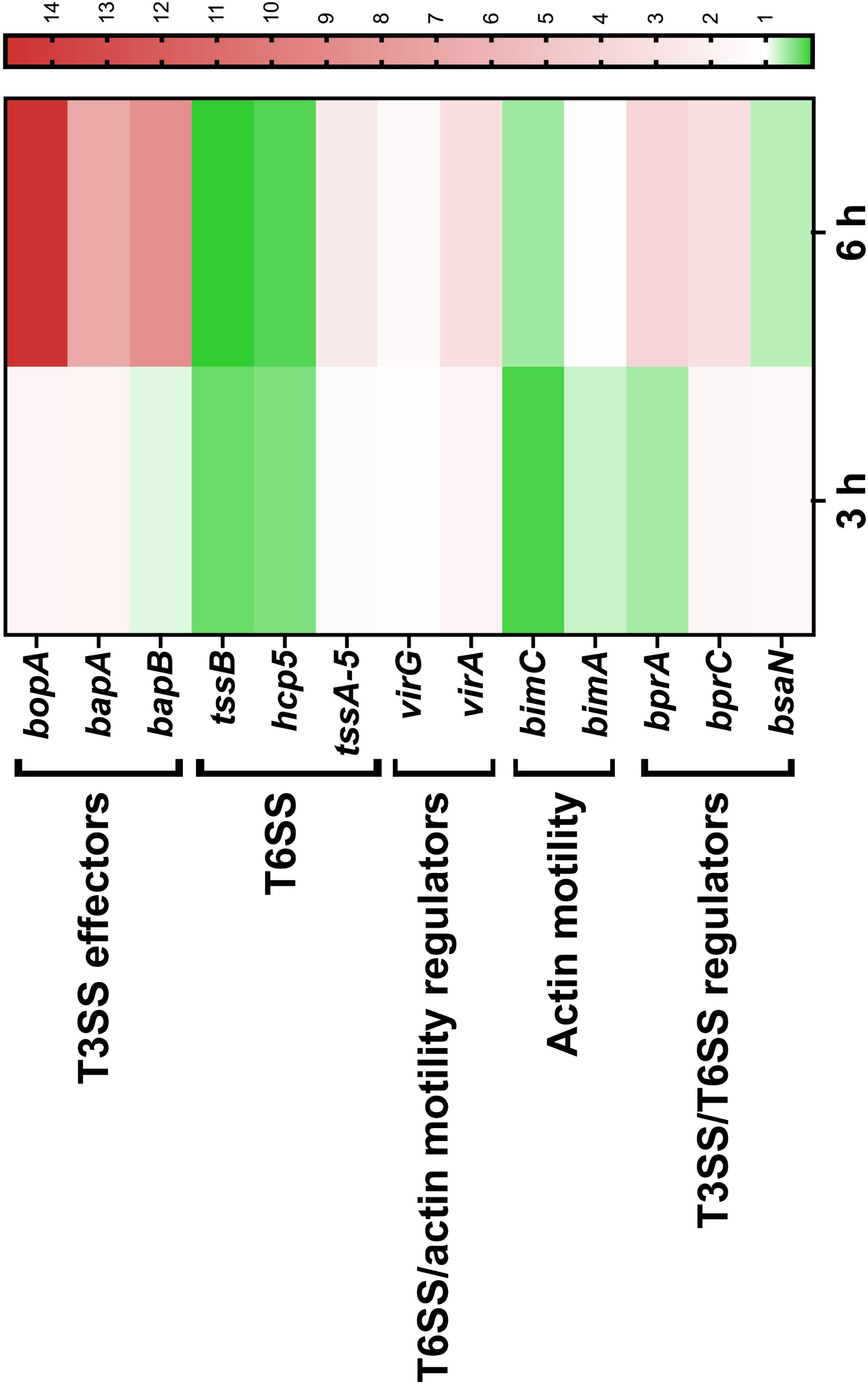
The *ΔbicA* strain displays disrupted expression of virulence factors during intracellular infection. RAW 264.7 cells were infected at an MOI of 10 with *Bpm* K96243 WT, Δ*bicA,* or Δ*bicA::bicA* for 3 or 6 h before being lysed and intracellular bacteria collected via differential centrifugation. Total bacterial RNA was collected from two independent experiments and used as a template for cDNA synthesis. Gene expression was measured by qPCR and the relative gene expression evaluated using the 2^-ΔΔCt^ method using WT as a control. The heatmap shows expression of Δ*bicA* relative to WT, meaning anything on the red spectrum is upregulated in Δ*bicA* and green is repressed in Δ*bicA*.

### The Δ*bicA* mutant is attenuated in an intranasal challenge model and results in differential macrophage recruitment and activation

Previously, we examined the virulence of Δ*bicA* in both acute and chronic models of gastrointestinal infection (1) and found that although Δ*bicA* demonstrated attenuation, those infection models are not able to discern the specific role of macrophages during infection. To fully investigate the virulence of Δ*bicA,* we intranasally challenged BALB/c mice with 3-5 LD_50_ *Bpm* K96243, Δ*bicA,* or Δ*bicA::bicA* and monitored their survival for 21 days (**Fig 4A**). All animals challenged with WT K96243 and Δ*bicA::bicA* succumbed to infection or reached the humane endpoint on day 4 post infection. Inversely, animals challenged with Δ*bicA* demonstrated 100% survival and showed no outward signs of infection besides minor body weight loss that was eventually recovered to near pre-challenge levels (**Fig 4A & B**). It should be noted that the body weight loss observed in a subset of Δ*bicA* challenged animals was delayed when compared to WT and Δ*bicA::bicA.* The reason behind the delayed onset weight loss is unclear but it could be related to the decreased intracellular replication phenotype seen *in vitro* (**Fig 1A & B**). The survivors of the Δ*bicA* challenge were euthanized 21 days post infection and lungs, liver, and spleen were collected for bacterial enumeration. Moderate amounts of bacteria were found in the lungs with very low titers in the liver and spleen (**Fig 4C**). One animal had high titers in both lungs and spleen and the spleen was enlarged with visible abscesses, this gross pathology is common in spleens chronically infected with WT *Bpm* (22). Once the attenuation of Δ*bicA* was established in this model, we sought to investigate the role of macrophages in during pulmonary infection. Another set of BALB/c mice were challenged with *Bpm* K96243, Δ*bicA,* or Δ*bicA::bicA* and at 48 h post challenge the lungs were harvested and divided to be processed for flow cytometric analysis and sectioned for pathological scoring. We designed a panel that allowed us to examine the macrophage populations within the lungs during infection and the basic activation state of those macrophages (Table 2). The gating strategy used to identify the total pulmonary macrophage populations in the lungs was modified from (23) but extra markers were added to assess the basic activation state of the populations identified. Macrophages have the ability to polarize to M1 and M2 subsets; M1 is the classic inflammatory profile associated with pathogen clearance while M2 subsets have immunoregulatory roles connected to tissue healing and limitation of inflammatory damage (20). Macrophage polarization is a complex system but for our purposes we have simplified the populations to M1-like expressing CD80 and CD86, while M2-like populations express CD163 and Arginase-1. These markers have allowed us to assess the basic activation states and gain insights into the dynamics of infection through the macrophage lens. Animals infected with WT K96243 and Δ*bicA::bicA* were found to have a higher total level of macrophages when compared to Δ*bicA* infected animals (**Fig 5A & B**). When those macrophage populations were examined for M1 and M2-like phenotypes we found that all groups had similar levels of M1-like activated macrophages but only WT and Δ*bicA::bicA* generated a small but distinctive M2-like populations (**Fig 5A & B**). Lung sections collected for pathological scoring were fixed in formalin, and H&E stained before being scored blind by a pathologist. Slides were scored on a 1-3+ system based on four criteria: nodules of inflammation, karyorrhectic debris/apoptosis, hemorrhage and congestion, and alveolar collapse. Pathology in the lungs of WT and Δ*bicA::bicA* infected animals was characterized by large nodules of inflammation that were densely populated by mononuclear cells and apoptotic debris with moderate amounts of congestion and alveolar collapse. Conversely, Δ*bicA* infected lungs were characterized by the complete lack of inflammatory nodules but slight to moderate congestion and hemorrhaging (**Fig 6A & B**). Interestingly, higher levels of infiltrating macrophages and M2-like populations found in WT and Δ*bicA::bicA* correlated with the presence of large inflammatory nodules and apoptotic debris but all conditions contain slight to moderate congestion and relatively equal M1-like populations. It is unclear whether this M2-like population elicited by the WT and Δ*bicA::bicA* is a response to uncontrolled bacterial replication and tissue damage or an intentional mechanism utilized by *Bpm*. Overall, these data demonstrate that Δ*bicA* is attenuated in an inhalational model and elicits a distinct pulmonary macrophage profile that correlates with pathological features disparate from WT and Δ*bicA::bicA*.

**Figure 4:**
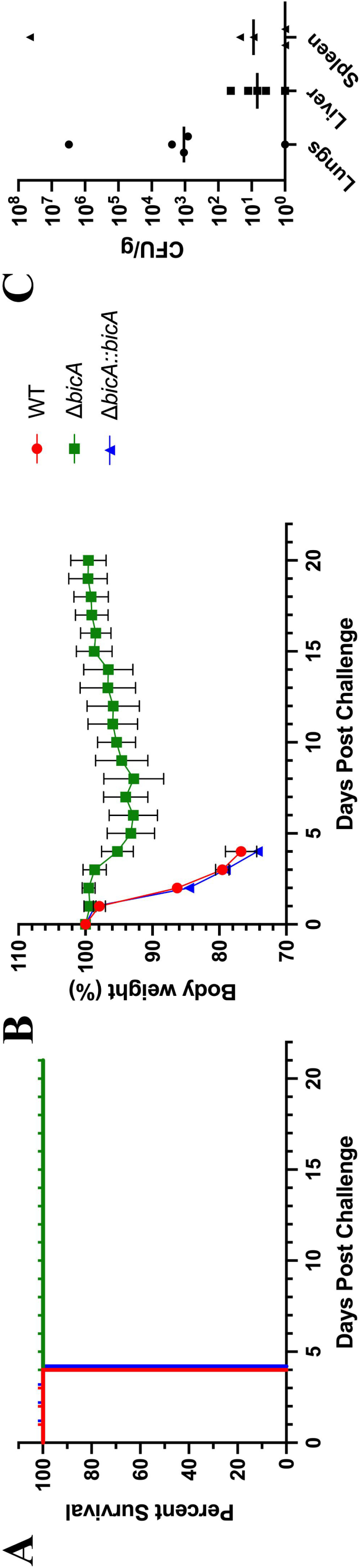
The *ΔbicA* strain is attenuated in the intranasal challenge model. BALB/c mice (n = 5/group) were intranasally challenged with 3-5 LD_50_ of *Bpm* K96243 WT, Δ*bicA,* or Δ*bicA::bicA* (1 LD_50_ ∼ 312 CFU) and monitored for 21 days post infection for survival (**A**) and weight loss (**B**). Animals were euthanized once the humane endpoint threshold was reached. On day 21 post infection, Δ*bicA* survivors were euthanized and lungs, liver, and spleen were homogenized for bacterial enumeration (**C**). Error bars in (**B**) represent SEM and lines in (**C**) represent median value.

**Table 2.**
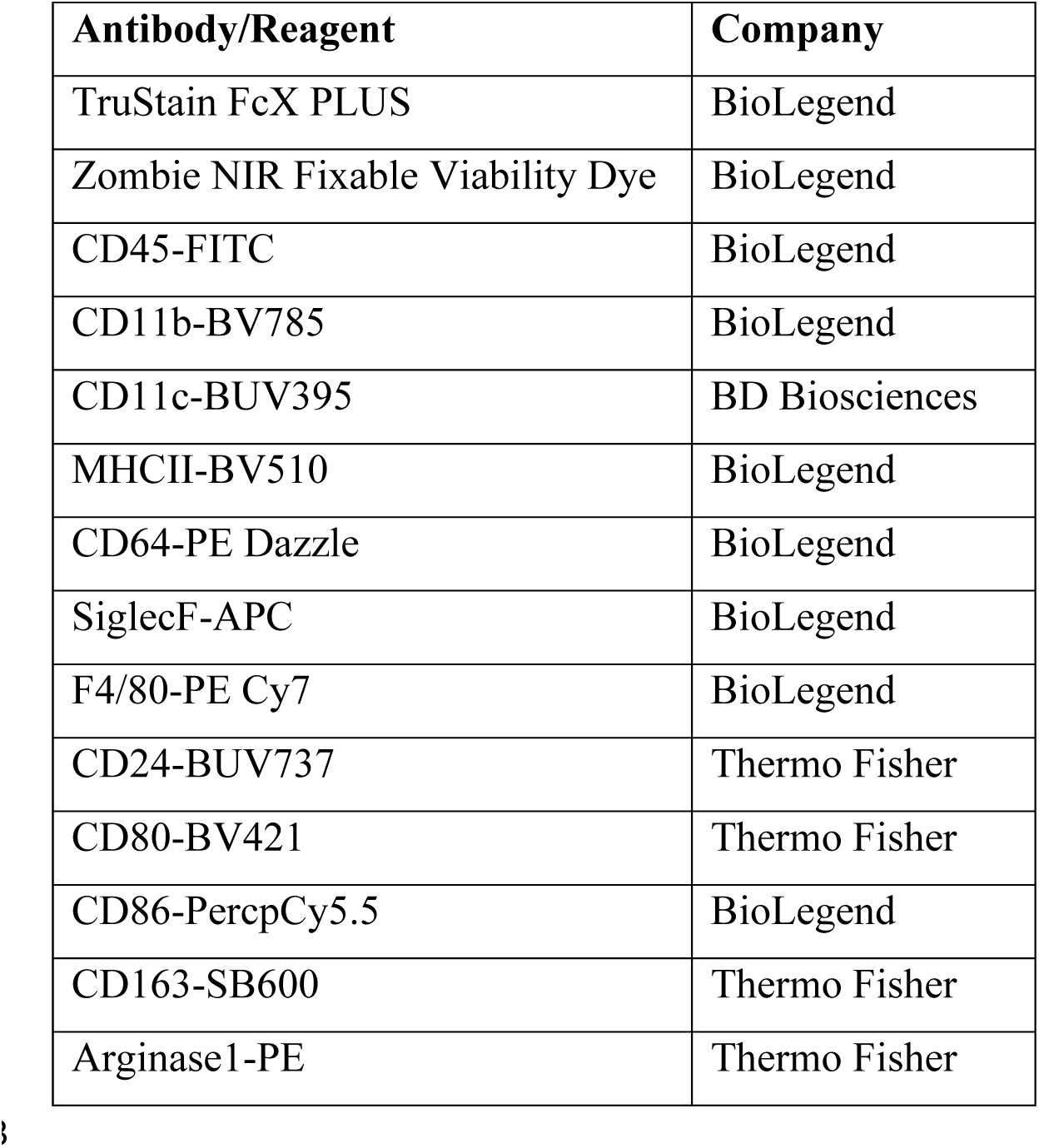
Flow cytometry antibodies and reagents

| Antibody/Reagent | Company |
| --- | --- |
| TruStain FcX PLUS | BioLegend |
| Zombie NIR Fixable Viability Dye | BioLegend |
| CD45-FITC | BioLegend |
| CD11b-BV785 | BioLegend |
| CD11c-BUV395 | BD Biosciences |
| MHCII-BV510 | BioLegend |
| CD64-PE Dazzle | BioLegend |
| SiglecF-APC | BioLegend |
| F4/80-PE Cy7 | BioLegend |
| CD24-BUV737 | Thermo Fisher |
| CD80-BV421 | Thermo Fisher |
| CD86-PercpCy5.5 | BioLegend |
| CD163-SB600 | Thermo Fisher |
| Arginase1-PE | Thermo Fisher |

**Figure 5:**
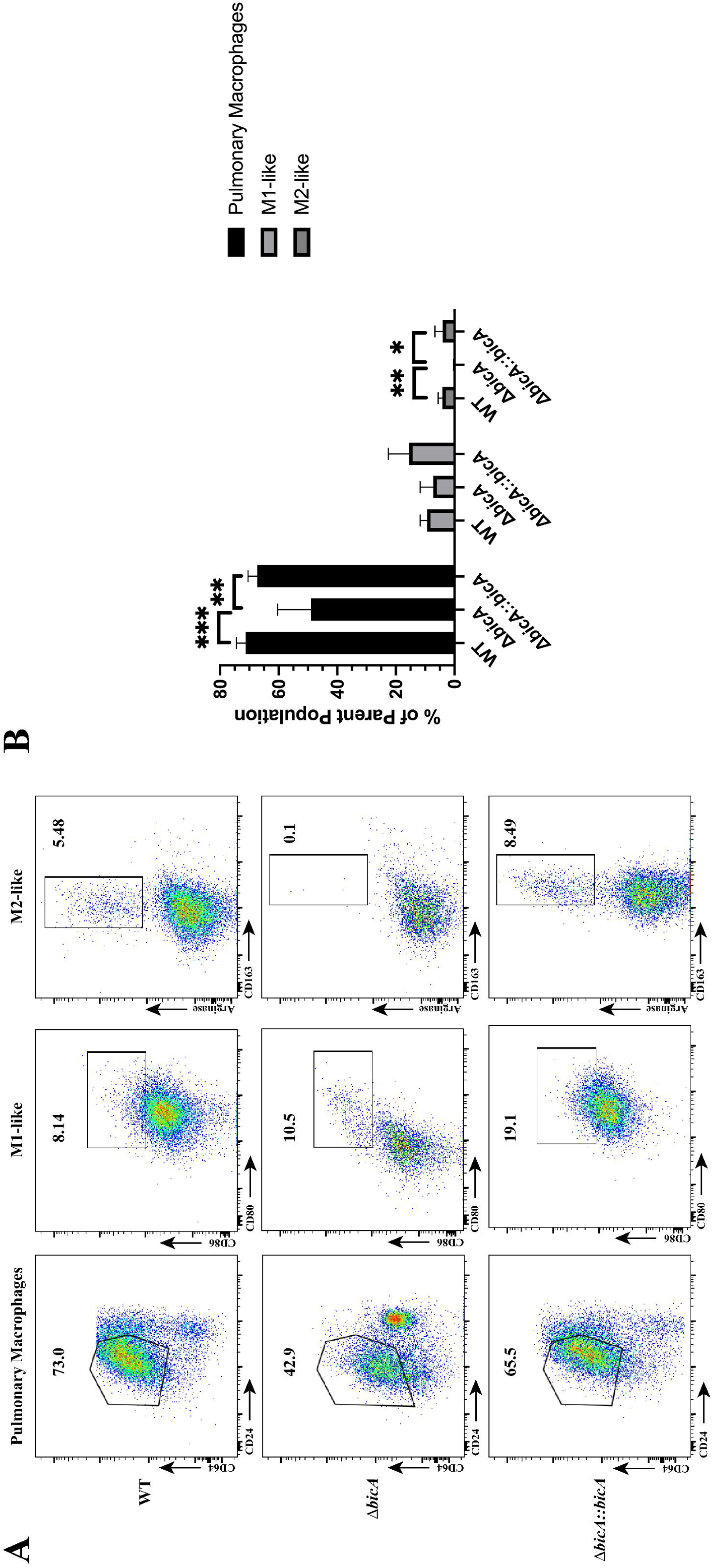
Different macrophage populations elicited by WT, Δ*bicA*, and Δ*bicA::bicA*. BALB/c mice (n = 5/group) were intranasally challenged with 3-5 LD_50_ of *Bpm* K96243 WT, Δ*bicA,* or Δ*bicA::bicA* (1 LD_50_ ∼ 312 CFU) and at 48 h post-infection lungs were harvested and processed for flow cytometry. A comprehensive gating strategy was adapted from (23) and activation markers were added to assess M1 (CD86 & CD80) and M2 (Arginase & CD163) polarization. Representative plots for pulmonary macrophages, M1-like, and M2-like populations (**A**) and populations from all animals (**B**) are shown. Bars represent mean ± SD and significant differences were assessed via one-way ANOVA followed by Sidak’s multiple comparison test. P < 0.05 *, p < 0.01**, p < 0.005***.

**Figure 6:**
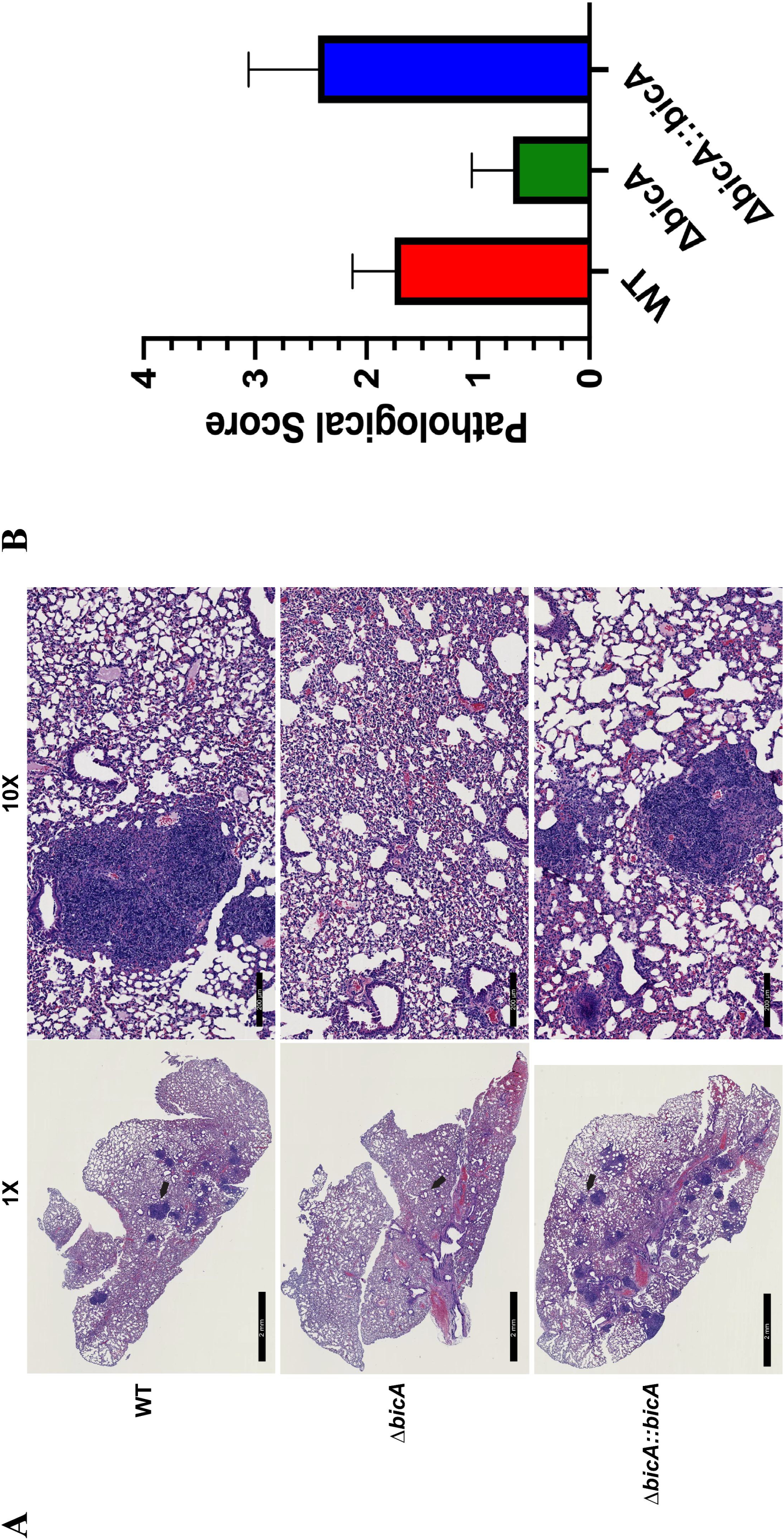
Distinct pathological features present in WT and Δ*bicA::bicA* but not Δ*bicA*. BALB/c mice (n = 5/group) were intranasally challenged with 3-5 LD_50_ of *Bpm* K96243 WT, Δ*bicA,* or Δ*bicA::bicA* (1 LD_50_ ∼ 312 CFU) and at 48 h post-infection lungs were harvested and portions were taken, formalin fixed, and H&E stained (n = 2/group). Representative images from WT, Δ*bicA,* and Δ*bicA::bicA* infected mice (**A)** 1x and 10x magnification. Lung pathology was scored using a 1-3+ system based on the following criteria: nodules of inflammation, karyorrhectic debris/apoptosis, hemorrhage and congestion, and alveolar collapse (**B**). Bars represent mean ± SD.

## Discussion

Melioidosis is a neglected tropical disease (24) that is a growing threat to nearly every continent on the globe. With recent findings demonstrating that *Bpm* is endemic to areas thought to be low burden or even non-endemic (6, 7, 25) and high incidence of antibacterial resistance, it is paramount that the underlying host-pathogen interactions are understood. The T6SS-mediated virulence is poorly characterized but previously, we performed dual RNA-seq on intestinal epithelial cells infected with WT or ΔT6SS *Bpm* to tease apart what bacterial and host factors are influenced by the T6SS. BicA was identified as differentially upregulated in the ΔT6SS mutant and then select for further characterization due to its implication in the T3SS and T6SS regulatory cascade. We found that a Δ*bicA* mutant was attenuated in both *in vitro* and *in vivo* models of gastrointestinal infection (1). To fully characterize the role of BicA during infection, we investigated its contribution during macrophage and pulmonary infection. As “professional phagocytes” macrophages bear the responsibility of being first responders to infection, pathogen uptake and clearance, antigen presentation, and immune modulation/coordination (26). The behavior of macrophages has a profound impact on the microenvironment and systemic recruitment of other immune cells. For these reasons macrophages are an attractive target for *Bpm,* as manipulation can skew the immune response to be advantageous for replication.

We chose two macrophage models, RAW 264.7 cells and BMDMs from BALB/c mice. RAW 264.7 cells are an immortalized cancerous cell line, so we wanted to compare any phenotype with primary cells from the same murine background to ensure that the phenotypes were not an artifact of these immortalized cells. The Δ*bicA* strain demonstrated an intracellular replication defect in both RAW 264.7 and BMDMs at 3, 6, and 12 h (**Fig 1A & B**). This defect was not due to a differential rate of phagocytosis (**Fig 1C**) but decreased intracellular survival starting as early as 1 h post-infection (**Fig 1D**). Δ*bicA* appeared to remain trapped within macrophages and did not form MNGCs (**Fig 2**) while the complemented strain recapitulated the WT features of actin motility and MNGC formation. Expression of some virulence factors was measured in Δ*bicA* and compared to WT at 3 and 6 h post-infection in RAW 264.7 cells (**Fig 3**) and it confirmed qualitative observations from **Fig 2**. Actin motility genes (*bimA* & *bimC*) and T6SS structural proteins (*hcp1* & *tssB*) were generally repressed in Δ*bicA* at 3 and 6 h; however, T3SS effector genes (*bopA, bopE, bapA,* & *bapB*) were upregulated at 6 h. Interestingly, there was activity on the *tssA-virAG* operon that was on par with WT but the downstream activity of T6SS genes *hcp1* and *tssB* was repressed. This suggests there might be a secondary signal for production of the T6SS that is lacking during Δ*bicA* intracellular infection. One possible explanation is that the lack of BicA is interrupting the co-activation/chaperoning of BsaN which feeds forward to activate the T6SS and actin motility. However, at 3 h *bsaN* and *bprC* expression was on par with WT which translated downstream to *virAG* activity even at 6 h when *bsaN* expression dips below WT levels. Downstream signaling events that should lead to T6SS activity occurred with diminished expression of *hcp1* and *tssB* so the lack of co-activator/chaperone activity is an unlikely source of the phenomenon. Alternatively, another explanation is that the Δ*bicA* strain lacks access to host cytoplasmic molecules that aid in expression of virulence factors. It has been shown that host glutathione aids in upregulating T6SS genes like *hcp1,* so it is possible that Δ*bicA* is being sequestered away from these molecules (27). The upregulation of T3SS effectors in Δ*bicA* could suggest that T3SS activity is upregulated, and the restrictive replication environment is potentially caused by increased expression T3SS proteins that are sensed by PRRs like NLR apoptosis inhibitory proteins (NAIPs) and results in inflammasome activation (28). However, the BicA homolog in *Salmonella,* SicA, is responsible for stabilizing or preventing degradation of SipB and SipC in the bacterial cytoplasm and Δ*sicA* has been shown to be partially complemented with the addition of *bicA* (29, 30). SipB and SipC form the translocon pore of the T3SS, so it is possible that Δ*bicA* is unable to produce an active T3SS. This notion is supported by reports that BicA is required for secretion of BopA and BopE (31). If this is the case, then Δ*bicA* is likely trapped in the phagosome for extended periods of time as the T3SS is responsible for escape, but mutants do exhibit an independent, albeit delayed, mechanism of escape (32). In **Fig 2**, Δ*bicA* appears primarily clustered at 3 and 6 h post-infection which might suggest they might be trapped in the phagosome as WT and Δ*bicA::bicA* replicate in a more diffuse pattern within the cytoplasm. Being trapped in the phagosome would restrict access to cytoplasmic host molecules like glutathione and at least partially explain the repression of the T6SS in the presence of regulatory events that should promote expression.

BicA has previously been implicated during inhalational melioidosis (33) but this was done with a transposon-based interruption of the gene so we sought to confirm the importance of BicA using our isogenic mutant, Δ*bicA*. When BALB/c mice were intranasally challenged with 3 - 5 LD_50_ of WT, Δ*bicA,* or Δ*bicA::bicA,* we observed one hundred percent survival in Δ*bicA* challenged animals, whereas all animals succumbed to infection on day 4 post-infection in the WT and Δ*bicA::bicA* groups (**Fig 4A**). The Δ*bicA* infected mice only presented a delayed but slight decrease in body weight that recovered to pre-challenge levels suggesting they were able to effectively control the infection (**Fig 4B**). On day 21, the Δ*bicA* survivors were euthanized and bacterial loads were assessed in the lungs, liver, and spleen. Low levels of bacteria were detected in all three organs with the lungs generally being the location of higher bacterial numbers (**Fig 4C**). It should be noted that all animals exhibited dissemination from the lungs to the liver and spleen but very few bacteria were recovered from these sites. One animal had robust replication in both lungs and spleen with visible abscesses on the organs, which is a feature that is common in WT infected organs, but this is likely an outlier event (22). The factors that created an environment conducive to Δ*bicA* replication in this animal are currently unknown.

Characterizing how intracellular pathogenesis events influence the host response is paramount to identifying avenues that can be targeted to combat the pathogen. To begin assessing this, we designed a flow cytometry panel to explore macrophage populations in the lungs during infection. We chose 48 h post-challenge to assess macrophage activity due to the stark contrast between WT/Δ*bicA::bicA* compared to Δ*bicA*; in our *in vivo* survival study at this timepoint the groups started to diverge in disease severity (**Fig 4C**). We reasoned that any differences at this timepoint would provide useful insight to the dynamics of the immune response to infection. A comprehensive gating strategy was devised using (23) as a guide with added polarization markers to delineate this portion of the inflammatory landscape. Although WT and Δ*bicA::bicA* infected animals elicited higher numbers of total pulmonary macrophages, all three experimental groups had equal levels of M1-like macrophages. Interestingly, WT and Δ*bicA::bicA* also had distinct M2-like populations (**Fig 5A & B**). This suggests that the presence of pro-inflammatory M1 macrophages aids in controlling bacterial replication but there is a larger, negative, contribution by M2 macrophages to promote replication. The phenomenon of skewing both M1 and M2 subsets is not unique to *Bpm,* it is a trait shared by *Mycobacterium tuberculosis, M. leprae,* and *Coxiella burnetti* (20). Both subsets can lead to downstream pathogenic effects on the host as uncontrolled inflammation from M1 can create excess tissue damage and conversely M2 can create an anti-inflammatory environment that allows pathogens to replicate undetected (20, 26). In conjunction with flow cytometry, we analyzed the tissue histopathology of a select number of mice in this cohort and the lungs were scored by a pathologist based on multiple criteria: nodules of inflammation, karyorrhectic debris/apoptosis, hemorrhage and congestion, and alveolar collapse. WT and Δ*bicA::bicA* infected lungs were characterized by large, discrete nodules of inflammation full of infiltrating mononuclear cells and apoptotic debris plus moderate amounts of congestion and alveolar collapse. The Δ*bicA* infected lungs lack the pronounced nodules but exhibit slight to moderate amounts of hemorrhage and congestion (**Fig 6A & B**). The nodules of mononuclear cells match the increase in pulmonary macrophages and are likely centered on replication hotspots. The presence of M2-like macrophages cannot be directly mapped to these foci but the centers being full of apoptotic debris increases the likelihood as M2 are more readily able to clear this debris through efferocytosis (34, 35).

In summary, we have explored the role of BicA in both immortalized and primary macrophages, demonstrating that Δ*bicA* has an intracellular survival defect. This defect is the result of a disruption in virulence factor expression defined by repression of the T6SS and actin motility; however, the cause of this repression is not fully understood. The Δ*bicA* mutant is highly attenuated in an inhalation model of melioidosis, inducing a differential macrophage recruitment and polarization profile paired with less severe histopathology. WT and Δ*bicA::bicA* infected mice recruit higher numbers of macrophages and promote a distinct M2 population that is absent in Δ*bicA,* suggesting that M2 polarization might be deleterious during infection.

## Materials and Methods

### Bacterial strains and growth conditions

All experiments were conducted with the prototypical wild-type strain of *Burkholderia pseudomallei* K96243 or derivative strains (K96243 *ΔbicA*, K96423 *ΔbicA::bicA*). K96243 *ΔbicA* and K96423 *ΔbicA::bicA* were constructed in (1). All strains were routinely grown at 37°C on LB agar plates and in LB broth with shaking.

### Macrophage culture conditions and infection assays

RAW 264.7 cells (ATCC TIB-71) were grown in Gibco Dulbecco’s Modified Eagle Medium (DMEM) plus 10% heat-inactivated fetal bovine serum (Gibco), 100 U/mL penicillin, and 100 μg/mL streptomycin (Gibco) at 37°C with 5% CO_2_. RAW 264.7 cells were maintained in T-75 flasks (Corning), detached using Accutase cell detachment solution (Biolegend) and seeded into 12 or 24 well plates (Corning). Bone marrow was collected from the femur and tibia of female BALB/c mice (Jackson Laboratories), RBCs lysed (Invitrogen 10x RBC Lysis buffer), and cells were added to polystyrene petri dishes (Sigma, 100mm x 20mm) containing RPMI 1640 w/ L-glutamine and HEPES (Gibco) plus 5 μM sodium pyruvate (Sigma), 100 U/mL penicillin, 100 μg/mL streptomycin (Gibco), 10% heat-inactivated fetal bovine serum (Gibco), and 25 ng/mL M-CSF (Biolegend). Cells were incubated at 37°C with 5% CO_2_ for 5 days with media changes on days 3 and 5. The resulting adherent cells were detached from the petri dishes using Accutase cell detachment solution (Biolegend) and seeded into 12 or 24 well plates (Corning) for further use.

RAW 264.7 cells or BMDMs were seeded at 5 × 10^5^/well in complete DMEM or RPMI without antibiotics into 24 well-plates and allowed to adhere overnight. *Bpm* strains were streaked on LB agar plates, grown at 37°C for 48 h, LB broth was inoculated and grown at 37°C with shaking for 12 h. Bacterial culture was diluted to 5 × 10^6^ CFU/mL in antibiotic free complete DMEM or RPMI and added to the cells for an MOI of 10. Cells were incubated with inoculum for 1 h for internalization, washed with PBS, and then media containing 1 mg/mL kanamycin was added for 1 h to kill off extracellular bacteria. For bacterial enumeration, cells were washed with PBS, lysed with 0.1% TritonX-100, serially diluted in PBS, and plated on LB agar plates. Percent intracellular replication was calculated by dividing the output at each timepoint by the presented input for each strain and converting the values to a percentage.

### Immunofluorescence assay

Infected BALB/c murine bone marrow-derived macrophage (BMDM) cells were fixed with 4% paraformaldehyde for 30 min following the select agent inactivation protocol approved by UTMB Environment Health and Safety. Cells were permeabilized with 0.25% Triton X-100 in PBS for 7 min at RT before incubation with serum (1:1000) from *Bpm* Δ*tonB*Δ*hcp1* (PBK001) live attenuated vaccine immunized mice (36). Cells were washed then incubated with 1:5,000 goat anti-mouse IgG, IgM (H+L) secondary antibody conjugated to Alexa 488 (Invitrogen) followed by actin and DNA staining using rhodamine phalloidin (Invitrogen) and DAPI (Sigma) together at 1: 10,000 dilutions. The coverslips were mounted using Prolong gold antifade (Molecular Probes, Life Technology) and sealed with nail polish. Stained cells were visualized using an Olympus BX51 upright fluorescence microscope and analyzed using ImageJ software from National institutes of Health (NIH).

### Measurement of gene expression by qRT-PCR

RNA for qRT-PCR analysis was prepared from two independent experiments of infected RAW 264.7 cells at 3- and 6-h post-infection at MOI 10. Bacterial RNA was isolated using Ambion TRIzol reagent (Life technologies) and Direct-zol RNA Miniprep kit (Zymo research). The cDNA was synthesized using the iScript cDNA synthesis kit (Bio-Rad) following the manufacturer’s protocol (Priming: 25°C, 5 min, reverse transcription: 42°C, 30 min, RT inactivation: 85°C, 5 min and store temperature: 4°C. The concentration and purity of cDNA were measured and normalized to 100 ng/ml for qRT-PCR step. The primers for qRT-PCR indicated in Table 1 were designed then tested for specificity by conventional PCR using Q5 High-fidelity DNA polymerase (New England Biolab). Gene expression was quantified using QuantiNova SYBR green (Qiagen) following the PCR cycling program as follow: initial heat activation step 95°C 2 min; 2-step 40 cycles of 5 s 95°C and 30 s 60°C. The threshold cycle (CT) and melting curve of each gene were automatically established and recorded by the software CFX Maestro Software (version 4.0). Relative gene expression level of each gene in Δ*bicA* mutant was normalized to wild-type strain using the 2^-ΔΔ^Ct method with 16S rRNA as reference gene.

### Intranasal challenge and survival studies

Female 6–8-week-old BALB/c mice (n = 5/group) (Jackson Laboratories) were intranasally (i.n.) challenged with 3-5 LD_50_ *Bpm* K96243, Δ*bicA,* or Δ*bicA::bicA* in 50 μL (25 μL/nare). One LD_50_ is equal to 312 CFU. Infected mice were monitored for survival and weight loss for 21 days post-infection and euthanized if the animal reached the threshold for humane endpoint. On day 21 post-infection, survivors were humanely euthanized, and lungs, liver, and spleen were collected for bacterial enumeration.

### Flow cytometry

Female 6–8-week-old BALB/c mice (n = 5/group) (Jackson Laboratories) were i.n. challenged with 3-5 LD_50_ *Bpm* K96243, Δ*bicA,* or Δ*bicA::bicA*; at 48 hpi, animals were euthanized, and lungs harvested for processing. Lung tissue was cut into small pieces and dissociated via incubation for 30 minutes at 37°C with slight rocking in RPMI plus 0.5 mg/mL collagenase IV and 30 μg/mL DNase I. The dissociated tissue was homogenized through a 100 μm cell strainer and fibroblasts and debris was pelleted via a 60 xg centrifugation for 1 minute. Supernatant was collected and RBCs were lysed for 5 minutes at RT. Following washes, pulmonary cells were adjusted to 1×10^6^ cells and stained using the reagents in Table 2. Briefly, cells were incubated with Zombie NIR (Biolegend) for 5 minutes in PBS, washed, and incubated with TruStain X plus (Biolegend) for 30 minutes followed by the extracellular antibodies (Table 2). Cells were fixed and permeabilized using Cytofix/Cytoperm (BD Biosciences) and stained for intracellular markers. Fully stained cells were resuspended in 4% ultrapure formaldehyde in PBS for 48 h in accordance with the inactivation protocol approved by UTMB Department of Biosafety before removal from BSL3 laboratory for analysis via BD Symphony full spectrum flow cytometer. Data were analyzed using FlowJo software. Proportions of parent populations were exported from FlowJo but, events within selected populations (pulmonary macrophages, M1-like, M2-like, etc.) were divided by total events selected from the previous parent gate.

### Evaluation of lung pathology

Lungs were collected from mice after humane euthanasia 48 h post-infection and fixed in 10% formalin for 48 h. Formalin fixed lung samples were submitted to the UTMB Anatomical Pathology core for paraffin embedding, mounting, and H&E staining. Stained slides were analyzed and scored by a pathologist (HLS) on a 1-3+ system. Slides were scanned and images taken using Aperio ImageScope.

### Statistical analysis

All statistical analysis was done using GraphPad Prism software (v9.0). P-values of < 0.05 are considered statistically significant. Survival differences were assessed via Kaplan-Meier survival curve followed by a log-rank test. An ordinary one-way ANOVA followed by Sidak’s multiple comparison test was used to analyze differences in intracellular replication and flow cytometry populations.

### Ethics statement

All manipulations of *Bpm* were conducted in CDC/USDA-approved biosafety level 3 (BSL3) laboratories at the University of Texas Medical Branch (UTMB) in accordance with approved BSL3 standard operating procedures. The animal studies were carried out humanely in strict accordance with the Guide for the Care and Use of Laboratory Animals by the National Institutes of Health. The IACUC protocol #0503014E was approved by the Animal Care and Use Committee of UTMB.

## Acknowledgements

This work was funded by USDA APHIS AP20VSD&B000C087, NIH NIAID grant AI12660101, and UTMB seed funds awarded to AGT. JLS is supported by a USDA APHIS NBAF Scientist Training Program Fellowship. We would like to thank Meredith Weglarz in the UTMB Flow Cytometry Core for the expertise and help in designing and implementing the flow cytometry experiments. We would also like to thank the lab of Janice and Mark Endsley for their expertise in lung dissociation and macrophages.

## Conflict of Interest Statement

The authors declare no conflicts of interest.

